# Effects of long-lasting insecticide net (LLINs) ownership/utilisation indicators on annual household malaria episodes (AHMEs) in three Health Districts in Cameroon

**DOI:** 10.1101/488445

**Authors:** Frederick Nchang Cho, Paulette Ngum Fru, Blessing Menyi Cho, Solange Fri Munguh, Patrick Kofon Jokwi, Yayah Emerencia Ngah, Celestina Neh Fru, Andrew N Tassang, Albert Same Ekobo

**Author notes:** Correspondence should be addressed to Frederick Nchang Cho.

## Abstract

**Introduction:** Household residents in malaria endemic areas are at high risk of multiple malaria episodes per year. This study investigated the annual household malaria episodes (AHMEs) in three health districts in Cameroon.

**Methods:** A community-based cross-sectional household survey using a multi-stage cluster design was conducted 2 – 3 years post campaign to assess long-lasting insecticide net (LLINs) ownership, utilisation, and maintenance as well as demographic characteristics. Multinomial regression analysis was used to identify factors associated with household LLIN ownership, utilization, and AHME.

**Results:** Household LLINs ownership, *de-facto* population with universal utilisation, and AHME were respectively, 92.5%, 16.0%, and 83.4%; thus, 4 out of 25 household residents effectively used LLINs the previous night. AHME was significantly (*p* < 0.05) associated with age and gender (OR; 1.6, 95% C.I; 1.1 – 2.3) of household head, health district (OR; 2.8, 95% C.I; 1.1 – 7.2) and tiredness (OR; 2.6, 95% C.I; 1.0 – 6.3). LLINs ownership and insufficiency also significantly contributed to AHME. The overall average cost for the treatment of malaria was 6,399.4±4,892.8Fcfa (11.1±8.5US$).

**Conclusions:** The proportion of households with at least one LLIN and those with at least one AHME were high. Findings are of concern given that average cost for the treatment of malaria represents a potentially high economic burden. The results outlined in this paper provide an important tool for the examination of the deficiencies in LLINs regular and universal utilisation.

## RÉSUMÉ

**Introduction:** Les maisons résidant dans les zones d’endémie du paludisme courent un risque élevé de subir plusieurs épisodes de paludisme par an. Cette étude a examiné les épisodes annuels de paludisme dans les maisons (EAPM) dans trois Districts Sanitaires du Cameroun.

**Méthodes:** Une enquête transversale de ménage basée sur la communauté utilisant une conception de grappe à plusieurs étapes a été conduite 2 - 3 ans après la campagne pour évaluer la possession, l’utilisation et l’entretien des moustiquaires imprégnées d’insecticide à longue durée (MILD) ainsi que les caractéristiques démographiques. Une analyse de régression multinomiale a été utilisée pour identifier les facteurs associés à l’appropriation, l’utilisation et l’entretien des MILD par les maisons.

**Résultats:** L’appropriation de MILDs par les maisons, la population *de-facto* avec l’utilisation universelle, et l’EAPM étaient respectivement, 92.5%, 16.0%, et 83.4%; donc, 4 sur 25 résidents des maisons ont effectivement utilisé des MILDs la nuit précédente. EAPM était significativement (*p* < 0.05) associé avec l’âge et le sexe (OR; 1.6, 95% C.I; 1.1 - 2.3) du chef de ménage, le district sanitaire (OR; 2.8, 95% C.I; 1.1 - 7.2) et la fatigue (OR; 2.6, 95% C.I; 1.0 - 6.3). La possession et l’insuffisance de MILD ont aussi contribué significativement à l’EAPM. Le coût moyen global pour le traitement du paludisme était de 6,399.4±4,892.8Fcfa (11.1±8.5US$).

**Conclusions:** La proportion des maisons avec au moins une MID et ceux avec au moins un EAPM étaient élevés. Les résultats sont préoccupants étant donné que le coût moyen pour le traitement du paludisme représente un fardeau économique potentiellement élevé. Les résultats décrits dans cet article fournissent un outil important pour l’examen des déficiences dans l’utilisation régulière et universelle des MILDs.

## 1. INTRODUCTION

Studies have identified the factors influencing the ownership and utilisation of long-lasting insecticide nets (LLINs) [1–11] in Cameroon and beyond. The utilisation rate of LLINs, especially amongst children less than five years old and pregnant women are widely low [2, 3, 12, 13], and is dropping with time from the date of ownership from a free mass distribution campaign (MDC) or purchase [14].

In malaria-endemic countries, especially those plagued with *Plasmodium falciparum* [15], malaria rates are still high, especially amongst the vulnerable population [16–18]. Malaria is a preventable and curable disease transmitted by the bites of the female Anopheles mosquitoes [16, 17, 19] and a serious global public health problem with an estimated 216 million cases in 91 countries in 2016 [17, 19–22]. Ninety per cent (90%) of all worldwide estimated malaria cases and 91% of deaths in 2016 occurred in 15 African countries alone contributing 80% of all cases [16, 20]. The epidemiology of malaria varies geographically depending on the local malaria transmission intensity or endemicity class as well as the *Plasmodium* species [15, 23]. The prevalence of malaria in Cameroon is 29% [24] and 15.0% in the North West and 46.1% in the South West Region amongst children under five in Cameroon [25].

The determinants of LLINs ownership, coverage, accessibility and utilisation are multiple and their contributions vary according to geographical location, sample size, and season of study [1, 8, 26–28]. Indicators of LLINs ownership and utilisation involve differences between health districts/localities, socio-demographic and economic statutes [12, 13, 29, 30].

The effective utilisation of LLINs has been reported to be invariably associated with ownership [4, 31], although annual household malaria episodes (AHME) is not primarily related to LLINs ownership. It is thought that poor LLINs utilisation by mostly the vulnerable is mostly due to behavioural attitudes of the population [6, 7, 13, 32], while the persistence of malaria is due in part to, underutilisation of LLINs, the use of other preventive methods, and negligence as well as vector resistance.

Studies in Cameroon and beyond have shown consistently that malaria is, and remains a public health problem [12, 22, 25, 33], with children in the ages of five or younger suffering between one-to-five or more malaria episodes per year [18]. Thus, in this study, the question is, “In health districts with high malaria endemicity and high LLINs ownership, what is the proportion and determinants of AHME, 2 – 3 years after the mass distribution campaign (MDC)?”.

## 2. MATERIALS AND METHODS

### 2.1 Free Mass Distribution Campaign

The Cameroonian Ministry of Public Health undertook a nationwide free LLIN distribution campaign from health facilities to all households in the country at the end of 2011, with the objective to provide an insecticide-treated net (ITN), with an optimal lifespan of 3 - 5 years [34], to all household beds or a LLIN for every two individuals per household, to a maximum of three ITNs per household, as described elsewhere [10, 12, 35].

### 2.2 Study area

The study was carried out in the BHD, SHD, and THD which constitute part of the most impoverished populations in Cameroon. These health districts are located in the North West and South West Regions of Cameroon. The characteristics of the study area have been described elsewhere [36].

### 2.3 Sampling design

This study is part of a prospective cross-sectional survey carried out in the THD in July and June 2017 and in Bamenda and Santa Health Districts in March to May 2018 [36].

### 2.4 Sample size determination

A minimum sample size of 385 for each health district was calculated as described elsewhere [36].

### 2.5 Recruitment procedures and measures

At enrolment, a structured questionnaire was used to record ownership of LLINs, utilisation of LLINs, and socio-demographic characteristics as well as housing and AHMEs.

### 2.6 Outcome variables

The main LLIN outcome variables were;

1. **LLINs ownership indicators**: *Ownership of LLINs, Coverage,* and *Access to LLINs within the household* were defined as described in other studies [10, 11, 37, 38].
2. **LLIN utilisation indicators**: *Household universal utilisation of LLINs* was defined as described in earlier studies [37–39]. *LLINs utilisation by the vulnerable population in the household* was defined as the proportion of children under five (or pregnant women) that slept under a LLIN the previous night [37]. *Regularly sleeping under bed nets:* household heads who reported habitually using nets daily [40]. *Slept under LLINs the previous night (SULPN)*: proportion of household heads who slept under a LLIN the previous night, as described in earlier studies [10, 11].
3. **Annual household malaria episodes (AHME)**: proportion of households which experienced at least one malaria episode in the last one year, where the numerator comprises the number of households surveyed wherein at least one household member suffered a malaria attack and the denominator, the total number of households surveyed.
4. **Independent variables** considered for association with LLIN ownership, use and AHME were age, gender, marital status, education, occupation, health district, house type, and household composition.

### 2.7 Statistical analysis

Data were analysed with IBM-SPSS Statistics 21.0 for windows (IBM-SPSS Corp., Chicago USA). The Chi square (χ^2^) test was used to compare socio-demographic characteristics with the AHME and multivariate logistic regression to identify significant correlates of the main outcomes. The level of statistical significance was set at *p* < 0.05.

## 3. RESULTS

### 3.1 Characteristics of study participants

A total of 1,251 household heads were surveyed, in the three health districts. The mean (±SD) age of study participants was 36.1 ±10.8, while the overall mean (±SD) household size was 4.7 ±2.1 members: 4.6 ±2.2 in BHD, 4.5 ±1.7 in SHD and 5.0 ±2.5 in THD. The overall mean AMHE was 2.2 ±1.7: 3.1 ±1.8 in BHD, 1.4 ±1.1 in SHD and 2.0 ±1.5 in THD. There was a significant association between AHME and house type as well as health district. Most (68.0%) households were headed by females, while majority (54.8 %) of the respondents were married. About 37.6% of the study participants had attained at least secondary school education and only 9.3% had no formal education (NFE). The greater percentage (35.3%) of the respondents was realised to be doing unskilled labour. Annual household malaria episodes (AHMEs) were frequent (89.2%) in households with surrounding bushes/farms or water pools (**Table 1**). Pregnant women were recorded in 93 (7.43%) of the households and children under the age of five in 766 (61.23%) of the households.

**Table 1:** Socio-demographic characteristics: by AHME.

| Independent variable | Dependent variable: |  | AHME |  |  |  |
| --- | --- | --- | --- | --- | --- | --- |
|  | Subclass | No | Yes | n (%) | p value | OR (95% C.I.) |
| Age groups (in years) | 20 | 13 | 17 | 30 (2.4) | < **0.001** | 0.2 (0.1 - 0.4) |
|  | 21 – 30 | 81 | 377 | 458 (36.6) | 0.233 | 0.7 (0.4 - 1.2) |
|  | 31 – 40 | 54 | 308 | 362 (28.9) | 0.620 | 0.9 (0.5 - 1.5) |
|  | 41 – 50 | 32 | 186 | 218 (17.4) | 0.935 | 1.0 (0.6 - 1.8) |
|  | 51 – 60 | 28 | 155 | 183 (14.6) | Ref | 1.0 |
|  | Mean age | 34.9±11.4 | 36.4±10.6 | 36.1±10.8 |  |  |
| Gender | Females | 135 | 716 | 851 (68.0) | 0.057 | 1.4 (1.0 - 2.0) |
|  | Males | 73 | 327 | 400 (32.0) | Ref | 1.0 |
| Marital status | Unmarried | 102 | 463 | 565 (45.2) | 0.464 | 1.1 (0.8 - 1.6) |
|  | Married | 106 | 580 | 686 (54.8) | Ref | 1.0 |
| Education | NFE | 13 | 103 | 116 (9.3) | 0.476 | 1.3 (0.6 - 2.6) |
|  | Primary | 62 | 308 | 370 (29.6) | 0.215 | 1.3 (0.8 - 2.1) |
|  | Secondary | 83 | 387 | 470 (37.6) | 0.624 | 1.1 (0.7 - 1.7) |
|  | Tertiary | 50 | 245 | 295 (23.6) | Ref | 1.0 |
| Occupation | Unemployed | 48 | 151 | 199 (15.9) | 0.728 | 1.2 (0.5 - 2.5) |
|  | Agricultural | 18 | 173 | 191 (15.3) | 0.337 | 0.7 (0.3 - 1.5) |
|  | Household & domestic | 6 | 49 | 55 (4.4) | 0.283 | 1.8 (0.6 - 5.7) |
|  | Unskilled | 79 | 362 | 441 (35.3) | 0.873 | 1.1 (0.5 - 2.2) |
|  | State/Parastatal | 43 | 173 | 216 (17.3) | 0.871 | 1.1 (0.5 - 2.3) |
|  | Professional | 14 | 135 | 149 (11.9) | Ref | 1.0 |
| House type | Caraboat | 10 | 74 | 84 (6.7) | 0.205 | 1.6 (0.8 - 3.4) |
|  | Mixed | 21 | 86 | 107 (8.6) | 0.286 | 1.3 (0.8 - 2.3) |
|  | Mud Block | 35 | 221 | 256 (20.5) | **0.043** | 1.6 (1.0 - 2.5) |
|  | Cement Block | 142 | 662 | 804 (64.3) | Ref | 1.0 |
| House size | 1 - 3 bedrooms | 190 | 951 | 1,141 (91.2) | 0.801 | 1.1 (0.6 - 1.9) |
|  | 4 - 7 bedrooms | 18 | 92 | 110 (8.8) | Ref | 1.0 |
|  | Mean number of bedrooms | 1.9±1.1 | 2.0±1.1 | 2.0±1.1 |  |  |
| Environmental factor | No | 21 | 114 | 135 (10.8) | 0.329 | 0.8 (0.4 - 1.3) |
|  | Yes | 187 | 929 | 1,116 (89.2) | Ref | 1.0 |
| Family size | 1 – 4 members | 103 | 519 | 622 (49.7) | 0.620 | 1.1 (0.7 - 2.0) |
|  | 5 – 7 members | 83 | 413 | 496 (39.6) | 0.768 | 1.1 (0.6 - 1.9) |
|  | ≥ 8 members | 22 | 111 | 133 (10.6) | Ref | 1.0 |
|  | Mean family size | 4.6±2.2 | 4.7±2.1 | 4.7±2.1 |  |  |
| Health District | Bamenda | 29 | 419 | 448 (35.8) | < **0.001** | 4.5 (2.5 - 8.2) |
|  | Santa | 98 | 287 | 385 (30.8) | **0.014** | 0.6 (0.4 - 0.9) |
|  | Tiko | 81 | 337 | 418 (33.4) | Ref | 1.0 |
|  | **Total** | 208 | 1,043 | 1,251 |  |  |

OR = Odds Ratio; C.I. = Confidence Interval; Ref = Reference group; **Boldface** numbers indicate significant *p* values

Of the 5,870 individuals (*de-facto* population) covered in the study, 4,908 (82.2%) spent the night in the 1,043 (83.4%) households which had suffered at least an AHME.

### 3.2 Ownership and utilisation of LLINs

A total of 2,958 LLINs were enumerated in the three health districts, overall LLINs density of 2.4 ±1.4. Long-lasting insecticide nets ownership, coverage and accessibility were 92.5%, 66.7% and 69.1% respectively. The utilisation rates were 14.6% for children less than five years old, 4.7% for expectant mothers and 16.0 % for entire households.

### 3.3 Determinants of household ownership and utilisation of LLINs

To investigate the determinants of LLINs ownership, coverage as well as utilisation in the three health districts, multinomial logistic regression was performed allowing adjustments for possible confounders. Households in the SHD (OR; 3.7, 95% C.I; 1.9 – 7.5, *p* <0.001) were significantly associated with LLINs ownership (**Table 3**). A majority of households with at least one LLIN (36.1%; 418/1,157) were found in the BHD, while (32.2%; 372/1,157) were in the THD. The difference was not statistically significant (*p* = 0.243). Secondary educational status, occupational status, and family size of 1 – 4 members were significantly (*p* >0.05) not associated with the ownership of at least one LLIN per household.

Being a household head in all the age groups except the 31 – 40 years old group, female, primary, and secondary educational level, BHD, and SHD and with no environmental factor were significant determinants associated with the use of LLINs by all children 0 – 5 years old in the household (**Table 3**). It is worth noting that the majority of the households with heads in the age group 21 – 30 years (35.4%; 184/520), females (68.7%; 357/520), secondary education (37.3%; 194/520) and BHD (48.1%; 250/520), had all children 0 – 5 years using LLINs compared with their contemporaries. Similarly, there was a significant association between household heads in the 21 – 30 years age group, BHD, families with sizes 1 – 4, and 5 – 7 members in the household and the use of LLINs by the entire household.

### 3.4 Annual household malaria episodes with LLINs ownership/utilisation indicators

A total of 4,908 (83.6%) of the 5,870 *de-facto* individuals were sampled in the 1,043 (83.4%) of households with at least one AHME in the last one year (**Table 2; Figure 1**). In terms of ownership indicators; AHMEs were associated with household accessibility (AOR; 1.2, 95% C.I; 0.6 – 2.5) to LLINs. Annual household malaria episodes were influenced by the use of LLINs by expectant mothers (AOR; 1.0, 95% C.I; 0.5 – 2.3), use of LLINs last night by the household head (AOR; 1.1, 95% C.I; 0.8 – 1.6), and regular utilisation of LLINs by the household head (AOR; 1.7, 95% C.I; 1.3 – 2.4), of which regular LLINs utilisation was significant (**Table 4**).

**Figure 1:**
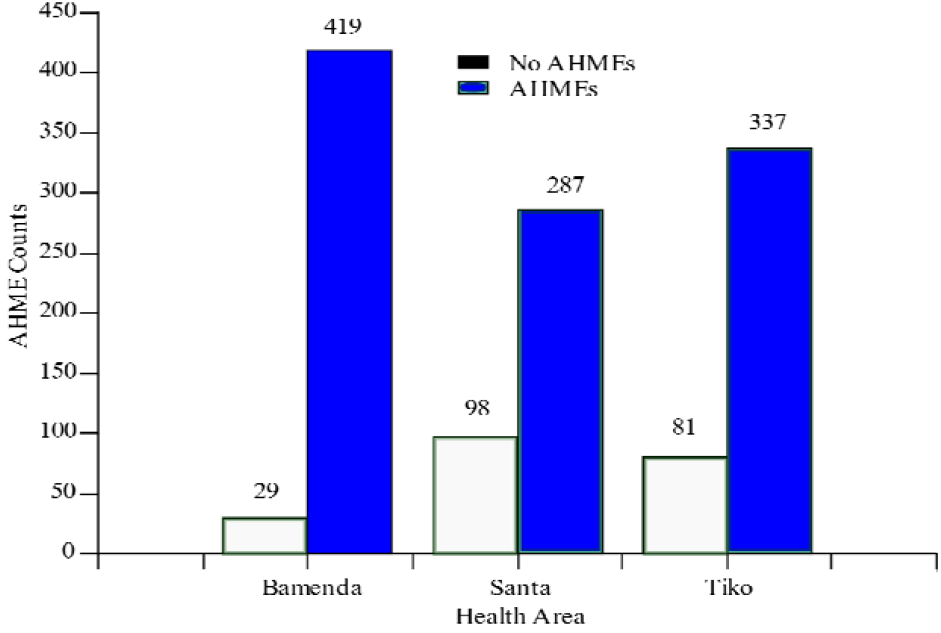
AHMEs by Health Districts

**Table 2:** Indicators of LLINs ownership/utilisation and AHMEs.

|  | Households |  |  |  |  |  | De-facto population in households |  |  |  |
| --- | --- | --- | --- | --- | --- | --- | --- | --- | --- | --- |
| Indicator | *n* (%) | BHD | SHD | THD | $\chi^2$ | *p* value | *n* (%) | BHD | SHD | THD |
| **Ownership** |  |  |  |  |  |  |  |  |  |  |
| At least one LLIN | 1,157 (92.5) | 418 | 367 | 372 | 12.23 | **0.002** | 5,577 (95.0) | 2,000 | 1,680 | 1,897 |
| Coverage | 836 (66.8) | 387 | 214 | 235 | 120.46 | **< 0.001** | 3,913 (66.7) | 1,893 | 937 | 1,083 |
| Accessibility | 865 (69.1) | 374 | 214 | 277 | 77.97 | **< 0.001** | 4,058 (69.1) | 1,825 | 937 | 1,296 |
| **Utilisation by** |  |  |  |  |  |  |  |  |  |  |
| Entire household | 256 (20.5) | 193 | 4 | 59 | 238.94 | **< 0.001** | 942 (16.0) | 767 | 10 | 165 |
| Children 0- 5 years | 520 (41.6) | 250 | 103 | 167 | 381.58 | **< 0.001** | 859 (14.6) | 427 | 188 | 244 |
| Expectant mothers | 59 (4.7) | 32 | 15 | 12 | 9.61 | **0.008** | 273 (4.7) | 173 | 46 | 54 |
| Regular utilisation | 484 (38.7) | 87 | 203 | 194 | 112.62 | **< 0.001** | 1,296 (22.1) | 346 | 297 | 653 |
| Household head last night | 350 (28.0) | 152 | 94 | 104 | 12.29 | **0.002** | 705 (12.0) | 356 | 111 | 238 |
| Installation | 811 (64.8) | 275 | 235 | 301 | 14.21 | **0.001** | 4,017 (68.4) | 1,347 | 1,138 | 1,532 |
| AHME | 1,043 (83.4) | 419 | 287 | 337 | 57.24 | **< 0.001** | 4,908 (83.6) | 1,900 | 1,290 | 1,718 |
| Mean AHME | 2.2±1.7 | 3.1±1.8 | 1.4±1.1 | 2.0±1.5 |  |  |  |  |  |  |

**Table 3:** Multinomial logistic regression of socio-demographic determinants of LLINs ownership and use by all children < 5 and entire household.

| Dependent variable: | Ownership of at least one LLIN |  |  | Used by children < 5 years old |  |  | Used by entire household |  |  |
| --- | --- | --- | --- | --- | --- | --- | --- | --- | --- |
|  | *n* (%) | *p* value | OR (95% C.I.) | *n* (%) | *p* value | OR (95% C.I.) | *n* (%) | *p* value | OR (95% C.I.) |
| Independent variable | *n* = 1,157 |  |  | *n* = 520 |  |  | *n* = 256 |  |  |
| **Age groups (in years)** |  |  |  |  |  |  |  |  |  |
| 20 |  |  |  | 16 (3.1) | **0.048** | 3.0 (1.0 - 8.6) | 6 (2.3) | **0.197** | 2.3 (0.7 - 7.8) |
| 21 – 30 |  |  |  | 184 (35.4) | **0.021** | 1.8 (1.1 - 3.0) | 111 (43.4) | **0.003** | 2.5 (1.4 - 4.7) |
| 31 – 40 |  |  |  | 146 (28.1) | 0.306 | 1.3 (0.8 - 2.2) | 76 (29.7) | 0.133 | 1.6 (0.9 - 3.0) |
| 41 – 50 |  |  |  | 108 (20.8) | **0.003** | 2.4 (1.3 - 4.2) | 35 (13.7) | 0.141 | 1.7 (0.8 - 3.4) |
| 51 – 60 |  |  |  | 66 (12.7) | Ref | 1.0 | 28 (10.9) | Ref | 1.0 |
| **Gender** |  |  |  |  |  |  |  |  |  |
| Female | 786 (67.9) | 0.751 | 0.9 (0.6 - 1.5) | 357 (68.7) | **0.008** | 1.6 (1.1 - 2.3) | 166 (64.8) | 0.583 | 0.9 (0.6 - 1.3) |
| Male | 371 (32.1) | Ref | 1.0 | 163 (31.3) | Ref | 1.0 | 90 (35.2) | Ref | 1.0 |
| **Marital status** |  |  |  |  |  |  |  |  |  |
| Unmarried | 512 (44.3) | 0.082 | 0.6 (0.4 - 1.1) | 176 (33.9) | **0.010** | 0.6 (0.5 - 0.9) | 98 (38.3) | **0.005** | 0.6 (0.4 - 0.8) |
| Married | 645 (55.7) | Ref | 1.0 | 344 (66.2) | Ref | 1.0 | 158 (61.7) | Ref | 1.0 |
| **Education** |  |  |  |  |  |  |  |  |  |
| NFE | 108 (9.3) | 0.577 | 0.8 (0.3 - 2.0) | 53 (10.2) | 0.576 | 0.8 (0.4 - 1.6) | 34 (13.3) | 0.291 | 1.4 (0.7 - 2.7) |
| Primary | 346 (29.9) | 0.814 | 0.9 (0.4 - 2.0) | 165 (31.7) | **0.002** | 2.1 (1.3 - 3.4) | 60 (23.4) | 0.973 | 1.0 (0.6 - 1.7) |
| Secondary | 421 (36.4) | **0.035** | 0.5 (0.2 - 1.0) | 194 (37.3) | **0.035** | 1.6 (1.0 - 2.4) | 95 (37.1) | 0.749 | 1.1 (0.7 - 1.7) |
| Tertiary | 282 (24.4) | Ref | 1.0 | 108 (20.8) | Ref | 1.0 | 67 (26.2) | Ref | 1.0 |
| **Occupation** |  |  |  |  |  |  |  |  |  |
| Unemployed | 182 (15.7) | **0.007** | 0.1 (0.0 - 0.6) |  |  |  |  |  |  |
| Agricultural | 174 (15.0) | **0.011** | 0.2 (0.1 - 0.7) |  |  |  |  |  |  |
| Household & domestic | 50 (4.3) | **0.035** | 0.2 (0.0 - 0.9) |  |  |  |  |  |  |
| Unskilled | 408 (35.3) | **0.014** | 0.2 (0.1 - 0.7) |  |  |  |  |  |  |
| State/Parastatal | 197 (17.0) | **0.014** | 0.2 (0.0 - 0.7) |  |  |  |  |  |  |
| Professional | 146 (12.6) | Ref | 1.0 |  |  |  |  |  |  |
| **Health District** |  |  |  |  |  |  |  |  |  |
| Bamenda | 418 (36.1) | 0.243 | 1.5 (0.8 - 3.1) | 250 (48.1) | **< 0.001** | 3.2 (1.9 - 5.3) | 193 (75.4) | **< 0.001** | 7.4 (4.2 - 13.0) |
| Santa | 367 (31.7) | **< 0.001** | 3.7 (1.9 - 7.5) | 103 (19.8) | **< 0.001** | 3.4 (2.0 - 5.9) | 4 (1.6) | **< 0.001** | 0.0 (0.0 - 0.1) |
| Tiko | 372 (32.2) | Ref | 1.0 | 167 (32.1) | Ref | 1.0 | 59 (23.1) | Ref | 1.0 |
| **House type** |  |  |  |  |  |  |  |  |  |
| Caraboat | 77 (6.7) | 0.357 | 1.5 (0.6 - 3.6) | 37 (7.1) | 0.518 | 0.8 (0.4 - 1.5) | 16 (6.3) | 0.92 | 1.0 (0.5 - 2.1) |
| Mixed | 103 (8.9) | 0.141 | 2.2 (0.8 - 6.5) | 35 (6.7) | **0.013** | 0.5 (0.3 - 0.9) | 7 (2.7) | 0.703 | 0.8 (0.3 - 2.1) |
| Mud Block | 239 (20.7) | 0.756 | 0.9 (0.5 - 1.7) | 115 (22.1) | 0.716 | 0.9 (0.6 - 1.4) | 58 (22.7) | 0.954 | 1.0 (0.6 - 1.6) |
| Cement Block | 738 (63.8) | Ref | 1.0 | 333 (64.1) | Ref | 1.0 | 175 (68.4) | Ref | 1.0 |
| **House size** |  |  |  |  |  |  |  |  |  |
| 1 - 3 bedrooms | 1,055 (91.2) | 0.96 | 1.0 (0.5 - 2.3) |  |  |  |  |  |  |
| 4 - 7 bedrooms | 102 (8.8) | Ref | 1.0 |  |  |  |  |  |  |
| **Family size** |  |  |  |  |  |  |  |  |  |
| 1 – 4 members | 551 (47.6) | **0.010** | 0.3 (0.1 - 0.7) | 113 (21.7) | **< 0.001** | 0.0 (0.0 - 0.1) | 175 (68.4) | **< 0.001** | 12.4 (6.0 - 25.7) |
| 5 – 7 members | 478 (41.3) | 0.912 | 0.9 (0.3 - 2.7) | 315 (60.6) | **0.007** | 0.4 (0.2 - 0.8) | 70 (27.3) | **0.007** | 2.7 (1.3 - 5.6) |
| ≥ 8 members | 128 (11.1) | Ref | 1.0 | 92 (17.7) | Ref | 1.0 | 11 (4.3) | Ref | 1.0 |
| **Own LLINs** |  |  |  |  |  |  |  |  |  |
| No |  |  |  | 2 (0.4) | **< 0.001** | 0.0 (0.0 - 0.2) | 1 (0.4) | **0.001** | 0.0 (0.0 - 0.2) |
| Yes |  |  |  | 518 (99.6) | Ref | 1.0 | 255 (99.6) | Ref | 1.0 |
| **Install LLINs beds** |  |  |  |  |  |  |  |  |  |
| No |  |  |  | 124 (23.9) | **< 0.001** | 0.5 (0.3 - 0.7) | 58 (22.7) | **< 0.001** | 0.4 (0.3 - 0.6) |
| Yes |  |  |  | 396 (76.1) | Ref | 1.0 | 198 (77.3) | Ref | 1.0 |
| **Environmental factor** |  |  |  |  |  |  |  |  |  |
| No |  |  |  | 64 (12.3) | **0.019** | 1.9 (1.1 - 3.3) | 46 (18.0) | 0.916 | 1.0 (0.6 - 1.7) |
| Yes |  |  |  | 456 (87.7) | Ref | 1.0 | 210 (82.0) | Ref | 1.0 |

**Table 4:**
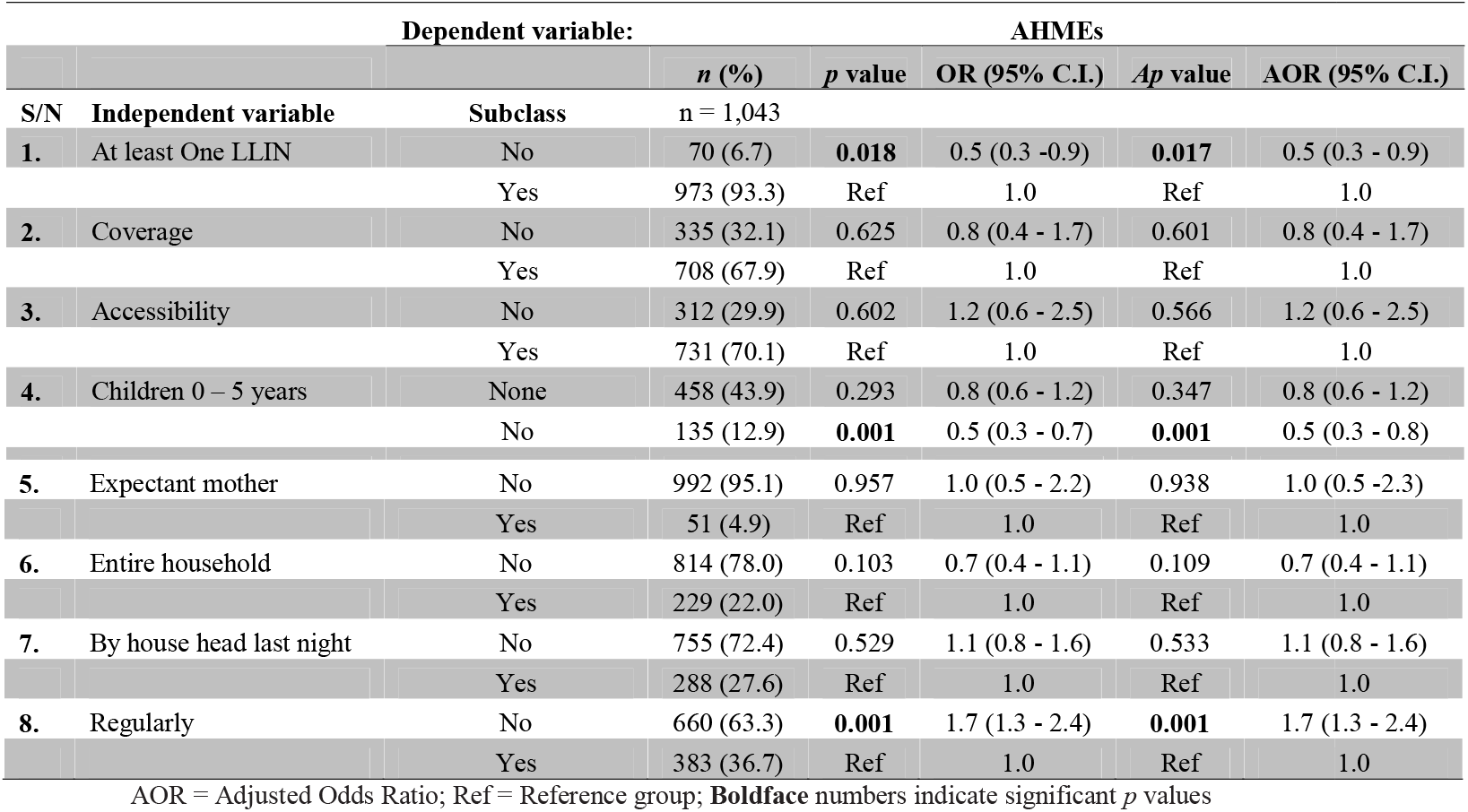
Multinomial logistic regression of LLINs ownership/utilization indicators in association with AHME.

### 3.5 Determinants of annual household malaria episodes

Annual household malaria episodes were associated with the age of the household head whereby households whose heads were 20 years old had the fewest AHMEs (*p* = 0.003) (**Table 5; Figure 2**). Multinomial analysis showed that the gender of the household head significantly (*p* = 0.017) influenced AHME. Households in the BHD had a higher AHME (*p* = 0.031) than those in the Santa and Tiko health districts (*p* > 0.05).

**Figure 2:**
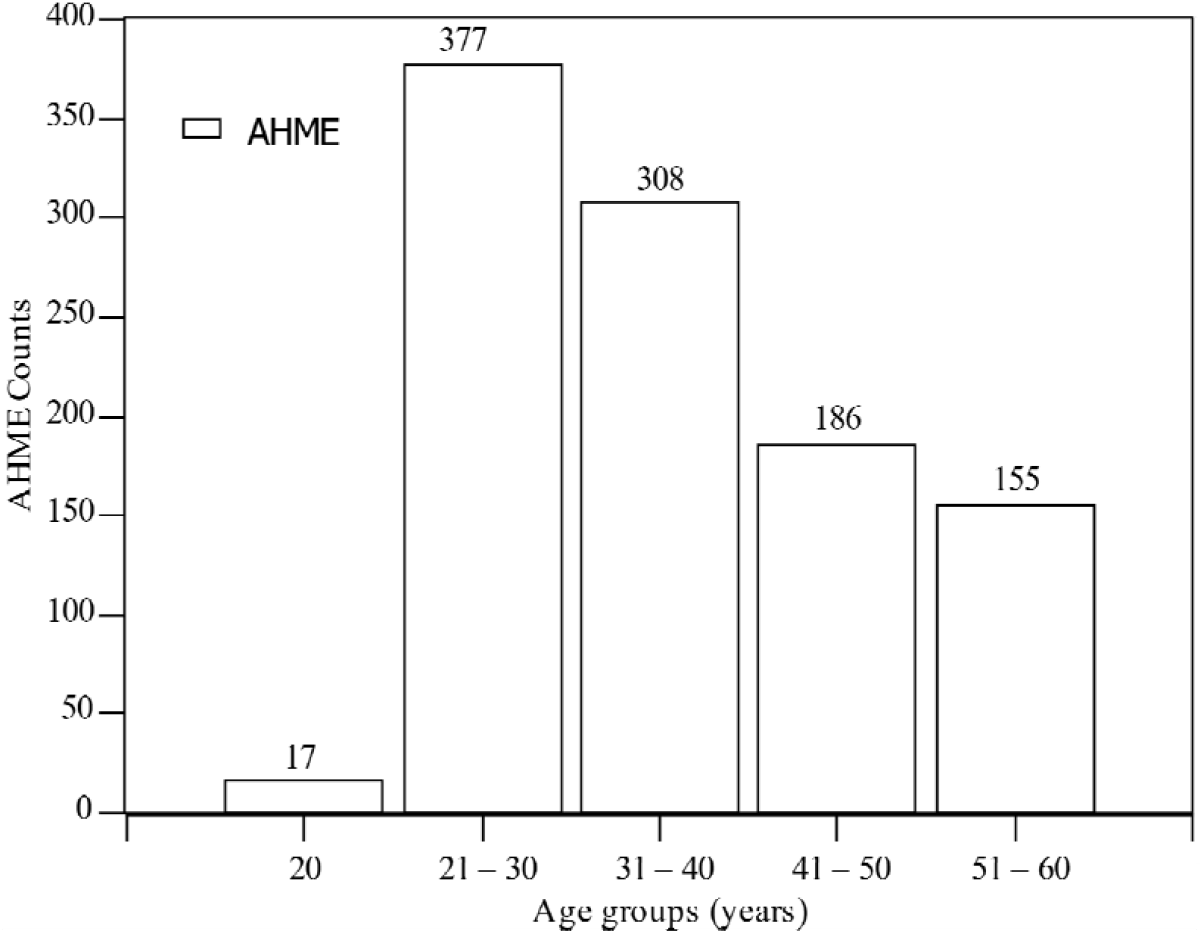
AHMEs by Age Groups

**Table 5:** Determinants of AHMEs.

| Dependent variable: |  |  | AHME |  |
| --- | --- | --- | --- | --- |
| Independent variable | Subclass | $n$ (%) | $p$ value | OR (95% C.I.) |
|  | 21 – 30 | 377 (36.1) | 0.332 | 0.8 (0.4 - 1.3) |
|  | 31 – 40 | 308 (29.5) | 0.649 | 0.9 (0.5 - 1.5) |
|  | 41 – 50 | 186 (17.8) | 0.846 | 1.1 (0.6 - 1.9) |
|  | 51 – 60 | 155 (14.9) | Ref | 1.0 |
| Gender | Female | 716 (68.6) | **0.017** | 1.6 (1.1 - 2.3) |
|  | Male | 400 (31.4) | Ref | 1.0 |
| Health District | Bamenda | 419 (40.2) | **0.031** | 2.8 (1.1 - 7.2) |
|  | Santa | 287 (27.5) | 0.251 | 0.7 (0.3 - 1.4) |
|  | Tiko | 337 (32.3) | Ref | 1.0 |
| Own LLINs | No | 70 (6.7) | **0.045** | 0.5 (0.2 - 1.0) |
|  | Yes | 973 (93.3) | Ref | 1.0 |
| Household LLINs sufficiency | No | 964 (92.4) | **0.002** | 0.1 (0.0 - 0.5) |
|  | Yes | 79 (7.6) | Ref | 1.0 |
| Tiredness | No | 975 (93.5) | **0.042** | 2.6 (1.0 - 6.3) |
|  | Yes | 68 (6.5) | Ref | 1.0 |

## 4. DISCUSSION

This study examined the possible causes of AHMEs in the Bamenda, Santa and Tiko Health Districts amidst high LLINs ownership, 2 – 3 years post nationwide free MDC. Overall, LLINs ownership was 92.5%, coverage was 66.8% (overall LLIN: Person ratio of 0.50) while the proportion of the *de-facto* population with universal utilisation was 16.0%, that of children less than five years old was 14.6% and AHMEs was experienced in 1,043 (83.4%) of the 1,251 households sampled. The overall average cost for the treatment of malaria was 6,399.4±4,892.8Fcfa (11.1±8.5US$): 9,010.3±5,297.2Fcfa (15.6±9.2US$) in BHD, 4,039.6±3,314.8Fcfa (7.0±5.7US$) in SHD and 5,774.5±4,325.1Fcfa (10.0±7.5US$) in THD.

### 4.1 Determinants of household LLINs ownership and utilisation

Long-lasting insecticidal nets (LLINs) ownership frequency is higher than 47 – 89.9% obtained elsewhere in Cameroon [3, 8, 12, 28], as well as 15.5 – 89.9% in Nigeria, Ethiopia and Myanmar [41–45] and in line with the 93.5% reported in Madagascar [46]. It was however, low compared to the 98.8% reported in Uganda [47]. The high frequency of LLINs ownership in this study could be attributed to the 2011 and 2015 free MDCs. However, the differences in LLINs ownership when compared with other studies, could be attributed to fluctuating ownership and universal coverage with time from the date of the MDC, and study designs [14, 44, 48], and study designs.

With respect to LLINs utilisation by the entire household, 16.0% of the *de-facto* population in 20.5% of the households and 14.6% of all the children less than five years in 41.6% of the households had at least used it the previous night. This low usage by the population is confirmed by other findings [41, 43, 46, 49] for the entire household and [8, 46, 47] for all children less than five years in the household. The very low levels of LLINs utilisation could be attributed to the fact that, even though the ownership of at least one LLIN per household was high; coverage was low, differences in the health districts, socio-demographic differences of the household heads, as well as the lack of sufficient space.

### 4.2 Annual household malaria episodes with LLINs ownership/utilisation indicators

The average cost for the treatment of uncomplicated malaria in Cameroon is 2,940Fcfa (6US$) [50]. The 83.4% AHMEs realised in this study is high compared to 50.8 – 77% reported in Nigeria, and Ghana [41, 49, 51, 52]. Associations were obtained between AHMEs and health districts (the BHD) as well as tiredness by the household head. The high AHMEs in this study is in line with a WHO report which states that the burden of malaria in low income countries is still high [16]. The AHME in this study is attributed to *Plasmodium falciparum* as opposed to *P*. *vivax* in Peru [53]. In this study, AHME was associated with LLINs accessibility as well as utilisation by expectant mothers, which is in line with a study reported in Zaria in Nigeria [49].

The average direct cost for the treatment of uncomplicated malaria in this study was 6,399.4Fcfa (11.1US$). This is low compared to the 65.1 US$ reported elsewhere in Cameroon [54], the 12.6 – 308 US$ reported elsewhere in Africa [51, 55–57], as well as 461.4 – 2,020.7US$ in the Slovak Republic [58]. It was however, in line with the 11.8 US$ reported in Vietnam [59] and higher than the 6US$ reported in Cameroon [50], 4.9 – 5.1US$ in Ghana and Ethiopia [60, 61]. The differences in the cost of the treatment of malaria might be due to, study designs, sample size and time of the study.

## 5. RECOMMENDATIONS

The Ministry of Health together with stakeholders should intensify education on the effective use of LLINs by all in the household, especially the vulnerable populations.

## 6. STRENGTHS AND LIMITATIONS OF THE STUDY

### 6.1 Strengths

The data used in this study was collected by trained surveyors, who had mastery of all the health areas in the study area. All the health district offices were consulted for the mapping of the health areas, quarters and census list of households used in the last MDC and Expanded Programme on Immunisation campaigns. The quality of data collected was assured through the multistage sampling strategy to minimize bias and pretesting of questionnaires.

### 6.2 Limitations

This was a cross sectional community-based study, carried out only in three health districts. Data was collected through self-reporting, with neither question on expenditure on malaria, nor one on diagnosis and type of malaria, rather, there was a question on the AHMEs.

In the calculation of the average expenditure on malaria, we did not distinguish simple from severe malaria.

## 7. CONCLUSIONS

In conclusion, the proportion of households with at least one LLIN and those with at least one AHME were high. The average cost for the treatment of malaria in the North and South West of Cameroon represents a potentially high economic burden, mainly to the Internally Displaced Persons and to the national economy as a whole. An implication is that increasing the universal utilisation could contribute to poverty reduction. The Ministry of Health, the National Malaria Control Program and other stakeholders need to identify mechanisms for ensuring that everybody has uninterrupted easy access to LLINs as well as regular utilisation.

## Data Availability

The dataset used for analysis in this study, was collected with the data collection tool as described in another study by PN Fru, FN Cho, AN Tassang, CN Fru, PN Fon and AS Ekobo [10], with DOI: https://doi.org/10.1155/2021/8848091 and cited in this study. A preprint with DOI: https://doi.org/10.1101/488445 of this study is also available online at www.biorxiv.org.

## ABBREVIATIONS

95% C.I: 95% Confidence Interval
AHME: Annual household malaria episodes
BHD: Bamenda Health District
LLINs: Long-lasting insecticide nets
MDC: Mass distribution campaign
NFE: No Formal Education
OR: Odds Ratio
*p*: Significance value
SD: Standard Deviation
SHD: Santa Health District
THD: Tiko Health District
χ^2^: Chi square

## DECLARATIONS

### Ethics approval and consent to participate

The study, obtained approval from the Institutional Review Board of the Faculty of Health Sciences, University of Buea. All participants gave oral/written consent prior to data collection.

### Authors’ contributions

Conceptualization and Methodology: FNC, PNF, and PNF. Field data collection: FNC, PNF, BMC, SFM, YEN, and PKJ. Data curation/Statistical analysis: FNC, PNF, and PKJ. Original draft preparation: FNC, PNF, BMC, SFM, PKJ, and PNF. Review and editing of draft: FNC, PNF, PKJ, PNF, and ASE. Administration and supervision: CNF, ANT, PNF, and ASE. All authors read and approved the final manuscript. Frederick Nchang Cho, Paulette Ngum Fru, and Patrick Kofon Jokwi contributed equally to this work.

## Acknowledgements

The authors are thankful to the heads of households who participated in this survey. We express immense gratitude to the community health workers and to the field assistants who worked under challenging field conditions to ensure that data was effectively collected.

